# Rationally Minimizing Natural Product Libraries Using Mass Spectrometry

**DOI:** 10.1101/2024.05.22.595232

**Authors:** Monica Ness, Thilini Peramuna, Karen L. Wendt, Jennifer E. Collins, Jarrod B. King, Raphaella Paes, Natalia Mojica Santos, Crystal Okeke, Cameron R. Miller, Debopam Chakrabarti, Robert H. Cichewicz, Laura-Isobel McCall

**Affiliations:** Department of Chemistry and Biochemistry, University of Oklahoma, Norman, Oklahoma 73019, United States; Department of Chemistry and Biochemistry, San Diego State University, San Diego, California, 92182, United States; Burnett School of Biomedical Sciences, University of Central Florida, Orlando, Florida, 32826, United States

**Author notes:** Authors contributed equally (Monica Ness, Thilini Peramuna).

## Abstract

Natural product libraries are crucial to drug development, but large libraries drastically increase the time and cost during initial high throughput screens. Here, we developed a method that leverages liquid chromatography-tandem mass spectrometry spectral similarity to dramatically reduce library size, with minimal bioactive loss. This method offers a broadly applicable strategy for accelerated drug discovery with cost reductions, which enable implementation in resource-limited settings.

## Main Text

Natural products play a major role in the development of novel pharmaceutical agents, accounting for nearly 70% of newly approved drugs in the past 40 years as direct natural molecules or as natural product mimics ^1^. Natural product discovery pipelines typically begin by screening large libraries of extracts (crude or pre-fractionated), then identifying and isolating bioactive candidates from these extracts. However, large libraries can result in long development times, high costs, and duplicate drug candidate identification (rediscovery). While efforts have been made to address these challenges, they suffer from several drawbacks ^2,3,4^. These methods do not consider MS/MS spectral similarity and did not directly evaluate applications of the minimalized libraries in the context of high throughput screening, or the retention of bioactive candidates. This latter data is critical to enable implementation within the broader drug discovery community, beyond natural product chemists. Here, we report the development of a new method to rationally reduce the size of natural product extract screening libraries by directly addressing cross-organismal redundancy in natural product production, through liquid chromatography-tandem mass spectrometry (LC-MS/MS) and molecular networking ^5^. This method led to (i) little loss of diversity; (ii) little loss of bioactive candidate molecules; (iii) increased bioactivity hit rate for a variety of whole-organism and purified protein targets; and (iv) greater library size reduction than previously published methods ^6^.

Our method uses an untargeted LC-MS/MS approach to choose extracts from a large natural product extract library. These extracts are complex and contain large numbers of different small molecules, some of which may be bioactive. Starting with MS/MS fragmentation patterns, data are then processed through GNPS Classical Molecular Networking software to group MS/MS spectra into scaffolds (**Fig. 1A**). These scaffolds are based on MS/MS fragmentation similarity, which correlates to structural similarity ^5^. Our rational libraries focus on scaffold diversity, as molecules with similar structures often demonstrate similar biological activity ^7,8,9^. Also, many drug development pipelines synthetically modify natural product scaffolds to improve structure-activity relationships ^10^. Therefore, diversifying core scaffolds, rather than individual molecules, can be prioritized in initial library design.

**Figure 1:**
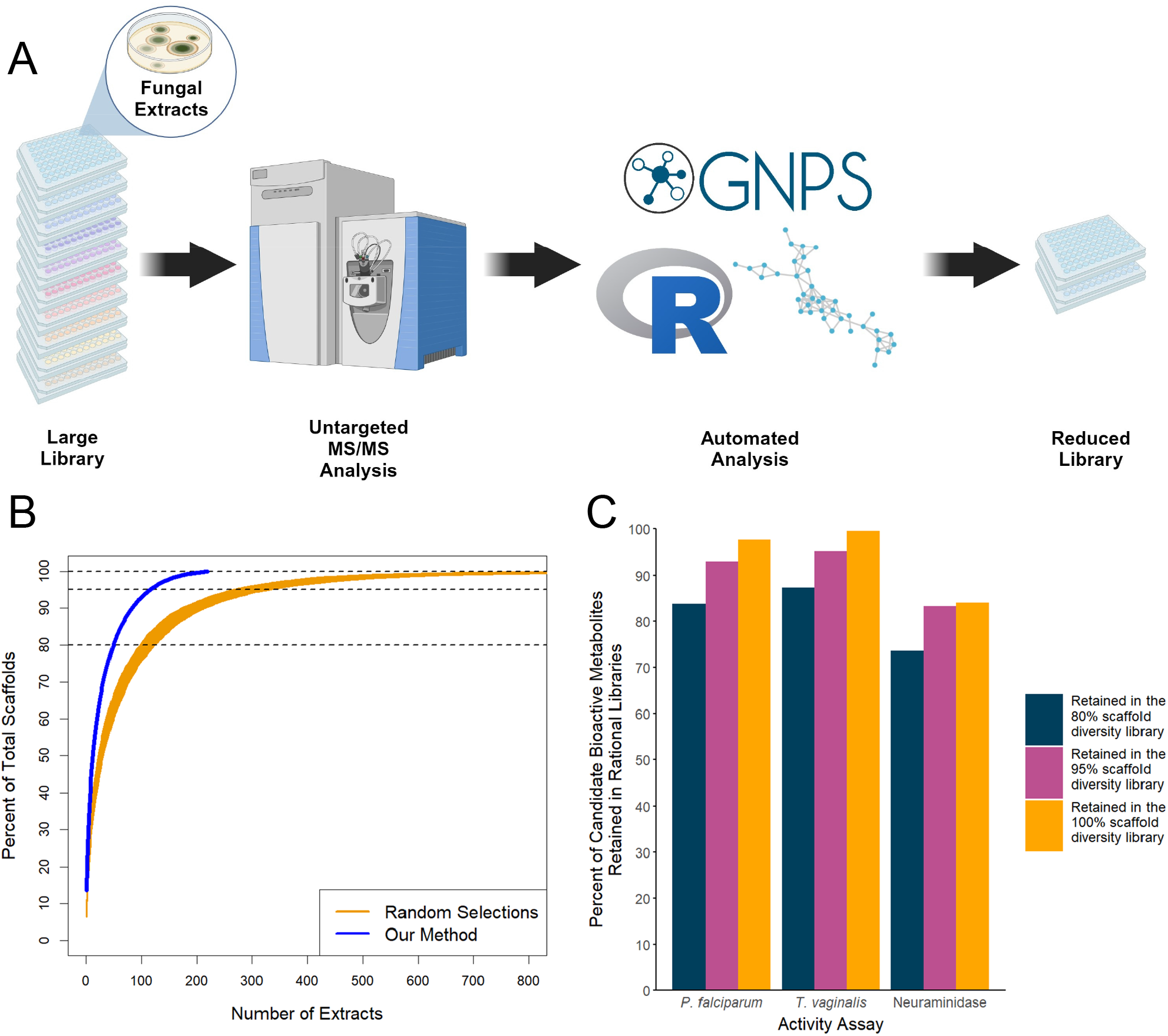
Rational library build using LC-MS/MS data decreases library size with limited loss of chemical diversity or bioactive candidate molecules. A) Method conceptual overview. B) Rapid aggregation of scaffold diversity using our method, outperforming random selections. C) High retention of bioactive candidate molecules in our rational libraries, for three diferent activity assays.

Using custom R code, which we make freely available (see Data Availability), our method selects the natural product extract with the greatest scaffold diversity. Next, the extract that contains the most scaffolds not already accounted for is added to the rational library. This process is iterated until a desired percentage of scaffold diversity is achieved in the rational library or a plateauing of diversity is observed. Compared to random selection, this accelerates the accumulation of diversity, and offers an 84.9% reduction in library size to reach maximal scaffold diversity. Specifically, in our evaluation library of 1439 fungal extracts, random sample selection achieved 80% of maximal scaffold diversity with an average of 109 extracts, whereas our method reached the same level of diversity with only 50 extracts. Similarly, to reach 100% scaffold diversity (representation in the library of all detected scaffolds), random selection required an average of 755 extracts. Our method only required 216 extracts (**Fig. 1B, Supplementary Table 1**). This size reduction exceeds the size reduction achievable with alternative methods ^6^.

Such dramatic library size reduction leads to concerns that key bioactive extracts will be lost. To assess bioactive extract loss in our rational libraries, we compared the bioactivity hit rate of the full library and of our minimal rationally-designed library on the eukaryotic parasites *Plasmodium falciparum* and *Trichomonas vaginalis*, as well as the influenza virus enzyme neuraminidase. Importantly, these assays represent two of the major types of assays in high throughput screening: phenotypic assays (*P. falciparum* and *T. vaginalis*), and target-based assays on purified enzymes (neuraminidase). To prevent bias, rational minimal library selection was blinded to bioactivity scores.

In the full library, the hit rate against *P. falciparum* was 11.3%, whereas the rational library designed to capture 80% scaffold diversity had an increased hit rate of 22%, and the rational library to 100% diversity had a hit rate of 15.7%. This pattern was replicated against *T. vaginalis* and neuraminidase (**Table 1**). Therefore, our method chooses extracts more likely to contain bioactivity, potentially because our method reduces the chemical redundancy inherent in natural product libraries ^2,11^. To confirm that these findings are not merely an artifact of the smaller library size, we compared our method with 1000 iterations selecting the same number of random extracts. In all cases, our method strongly outperformed random extract selection (**Table 1** and **Supplementary Table 2**)

**Table 1.** Higher hit rates for rational libraries.

| Activity assay | Hit rate in full library (1439 extracts) | Hit rate in the 80% scaffold diversity library (50 extracts) | Lower and upper quartile hit rates for 50 random extracts (1000 iterations) | Hit rate in the 95% scaffold diversity library (116 extracts) | Hit rate in the 100% scaffold diversity library (216 extracts) |
| --- | --- | --- | --- | --- | --- |
| *P. falciparum* | 11.26% | 22.00% | 8.00%-14.00% | 19.83% | 15.74% |
| *T. vaginalis* | 7.64% | 18.00% | 4.00%-10.00% | 17.24% | 12.50% |
| Neuraminidase | 2.57% | 8.00% | 0.00%-2.00% | 6.03% | 5.09% |

To further address concerns with regards to loss of bioactive molecules, we identified molecules correlated with bioactivity in the full library, and assessed whether they were retained in the minimal rational library. We found that, of the 266 molecules significantly correlated with anti-*Plasmodium* activity in the full library (r > 0, p >0.05, Bonferroni corrected), 223 were retained in the rational library at 80% scaffold diversity (84%), and 260 were retained in the 100% diversity library (98%, **Fig. 1C**). A similar conclusion was reached with regards to molecules correlated with anti-*T. vaginalis* and anti-neuraminidase activity (**Fig. 1C and Supplementary Table 3**).

To validate our method and demonstrate its utility beyond our laboratory, we applied the same rational library-building process to LC-MS data collected by independent investigators and tested for bioactivity against another pathogen, the parasite *Trypanosoma cruzi* ^12^. After rationally reducing their full library size (1600 total extracts) down to 104 extracts with our method, we also observed a rapid accumulation of chemical diversity, an increased hit rate in the rational libraries when compared to the full library (from 0.5% to 1.92%), and the retention of 78-95% of features correlated with bioactivity (See **Supplementary Tables 4-5, Supplementary Fig. 1-2**).

In addition to the accelerated screening and cost reduction enabled by our rational library development method, there are also many other advantages of the LC-MS/MS data acquired through this pipeline. Any bioactive candidates identified in a screen of these rational libraries will already have LC-MS/MS data readily available for initial structure elucidation. This data can also be input into many convenient platforms for dereplication ^13,14,15,16^ and structural annotation ^5,17,18^. Abundance information from LC-MS data can also be leveraged to build bioactive correlations ^19^ and identify candidate bioactive molecules, as demonstrated in this study. Such data can also be used to guide compound isolation and purification for definitive structure elucidation by NMR. If desired, extracts containing molecules previously described as promiscuously toxic, or bioactive in the target biological system can also be excluded, to prevent re-discovery. Alternatively, in under-studied diseases, investigators may wish to retain these latter extracts, to enable repurposing of these known compounds to other disease models.

The results presented above focus on positive mode data, as more molecules can be detected in positive mode ^20^. However, we found the same conclusions when analyzing negative mode data (**Supplementary Tables 7-8, Supplementary Fig. 3-4)**. Likewise, modulating GNPS Classical Molecular Networking parameters ^5^ or mirroring low-resolution data acquisition parameters had limited impact (**Supplementary Tables 9-11, Supplementary Fig. 5-6, Supplementary Data 1)**. These observations indicate a broad tolerance of our approach to data acquisition and processing parameters.

While library screening costs represent a small proportion of the total drug development investment, they remain a preliminary bottleneck that is especially critical in settings where financial resources are limited, such as neglected tropical diseases or rare conditions. By reducing the size of the library to be screened using our method, additional screening, dose-response assessment, and counter-screening assays can be implemented, leading to reduced false positive rates. Rationally-designed library size reduction also enables the implementation of more complex (and thus more costly per-well) assays that better mimic *in vivo* conditions, increasing translatability and likelihood of clinical success. While we acknowledge the initial up-front cost and time needed for the LC-MS analysis, this is mitigated by the fact that it can enable cost reduction across all subsequent high throughput screening projects using this library (as demonstrated here), prevent re-discovery of known bioactives, and expedite bioactive candidate structure elucidation. This method should also be suitable for libraries of pre-fractions, or to prioritize crude extracts for fractionation, leading to further cost reduction and time savings. Lastly, our method was applicable to datasets and screening assays implemented outside our laboratory, demonstrating its broad utility.

## Methods

### Collection of Fungal Extracts

The University of Oklahoma’s Citizen Science Soil Collection Program ^21^ asked citizen scientists to submit soil samples that were then used to isolate fungi. These fungi were grown to generate extracts that are used in various drug discovery efforts. Fungi were selected for the library based on morphological analysis, and the unique fungi from each soil sample were retained. ITS-based taxonomic identifications were also generated. Over 79,000 fungi originating from all fifty states and the District of Columbia are included in the fungal library (<u>Citizen Science Soil Collection Program (shareok.org)</u>). The subset of fungal extracts used in this study were isolated from 443 soil samples with a broad distribution across the United States (**Supplementary Fig. 7**) and includes members of 141 different genera (**Supplementary Data 2**). Taxonomic identification was accomplished via sequencing of the internal transcribed spacer region of the genomic DNA ^3^. These fungal isolates were cultured for three weeks on a solid-state medium composed of Cheerios breakfast cereal supplemented with a 0.3% sucrose solution containing 0.005% chloramphenicol, as previously optimized ^22, 23^.

### Metabolite sample preparation

Samples for bioactivity assays and LC-MS analysis were prepared on an automated platform that combined both extraction and partitioning steps as previously described ^24^. Fungal cultures prepared in 16-by 100-mm borosilicate tubes were subjected twice to a vol:vol water:ethyl acetate partitioning. The organic solvent was removed *in vacuo* and the remaining organic residues were stored at -20°C until resuspended in DMSO for screening. These same extracts were diluted 1:10 in methanol containing sulfadimethoxine as an internal standard for liquid chromatography-tandem mass spectrometry (LC-MS/MS) analysis.

### *Plasmodium falciparum* Assay

Parasites were cultured following a protocol by Trager and Jensen, with minor modifications ^25, 26^. Specifically, the multidrug resistant *P. falciparum* line Dd2 was grown at 37°C in 5% CO_2_ with Rosewell Park Memorial Institute media (RPMI) 1640 media supplemented with 25 mM of 4-(2-hydroxyethyl)-1-piperazineethanesulfonic acid (HEPES) pH 7.4, 26 mM NaHCO_3_, 2% dextrose, 15 mg/L hypoxanthine, 25 mg/L gentamicin, and 0.5% Albumax II in human A+ blood. The impact of fungal extracts on *P. falciparum* growth was measured via a SYBR Green I fluorescence-based assay as described ^25^ . Extracts were resuspended in DMSO and plated on microtiter plates with asynchronous Dd2 culture at 1% parasitemia, 1% hematocrit, for a final concentration of 2 µg/mL, maintaining a DMSO concentration of < 0.25% in all cases to avoid assay interference. Culture plates were then incubated with extracts under standard growth conditions for 72 h. After incubation, assay plates were frozen at -80ºC and subsequently thawed to promote lysis. Once thawed, an equal volume of lysis buffer (20 mM Tris-HCl, 0.08% saponin, 5 mM ethylenediaminetetraacetic acid (EDTA), and 0.8% Triton X-100) with 1X SYBR Green I was added. Plates were incubated for 45 min - 1 h, protected from light, prior to fluorescence reading at excitation 485 nm, emission 530 nm, on a Synergy Neo2 multi-mode reader (BioTek, Winsooki, VT). Values were then normalized to 10 mM chloroquine and vehicle only control wells. Extracts showing more than 75% inhibition of parasite growth were considered hits. Bioactivity assays were performed separately from rational library generation; investigators performing this assay were unaware of the rationally-selected extracts, and vice versa.

### *Trichomonas vaginalis* Assay

*Trichomonas vaginalis* Donne (PRA-98) from the American Type Culture Collection (ATCC, Bethesda, MD) was grown at 37°C in filter sterilized Keister’s Modified TYI-S33 medium (2% (w/v) casein, 1% (w/v) yeast extract, 55.6 mM glucose, 34.2 mM NaCl, 4.4 mM KH_2_PO_4_, 5.7 mM K_2_HPO_4_, 12.7 mM l-cysteine HCl, 1.1 mM l-ascorbic acid, 86.7 μM ferric ammonium citrate, 10% (v/v) heat inactivated fetal bovine serum, 0.052% (w/v) bovine bile salts in 1 L of Millipore water). Micro aerophilic conditions were maintained with the use of BD GasPak EZ Campy sachets. Samples consisting of 4 × 10^4^ trichomonads per well were treated with DMSO, 25 μM metronidazole, or fungal extracts at 10 μg/mL, not exceeding 0.5% DMSO. After a 16 h incubation, cells were fixed (1% glutaraldehyde, 5 μM propidium iodide, and 5 μM acridine orange in PBS). After a 3 h incubation at 37 °C, cells were imaged using a PerkinElmer Operetta high-content imaging system. Data were analyzed using the Harmony 3.5.1 software package as previously described ^22^. Number of live cells imaged was normalized to the DMSO control (100% growth). Extracts were considered active if they inhibited growth of *T. vaginalis* by more than 80%. Bioactivity assay was performed separately from rational library generation; investigators performing this assay were unaware of the rationally-selected extracts, and vice versa.

### Neuraminidase Assay

Fungal extracts plus 0.01% TritonX to break up aggregates were tested in high throughput format at 10 µg/mL using a commercial neuraminidase activity assay, according to manufacturer instructions (Sigma Aldrich, catalog number MAK121). Those identified as reducing neuraminidase activity to less than 30% of vehicle control were retested in a dose-response curve to verify activity. Hits were considered confirmed if they showed dose dependent inhibition. Briefly, the reaction mixture containing buffer, substrate, cofactors, enzyme, and dye were mixed with the extracts and incubated at 37°C. Absorbance at 570 nm was recorded at 20 and 50 minutes. Activity of the neuraminidase enzyme was calculated according to the kit instructions. Activity scores were then normalized. Bioactivity assay was performed separately from rational library generation; investigators performing this assay were unaware of the rationally-selected extracts, and vice versa.

### LC-MS/MS Data Acquisition

Fungal extract separation was done with Thermo Scientific Vanquish ultra-high-performance liquid chromatography (UHPLC) instrument equipped with a Kinex 1.7 µm C18 50 x 2.1 mm LC column with a C18 guard cartridge (Phenomenex). The mobile phases used were water with 0.1% formic acid (mobile phase A), and acetonitrile with 0.1% formic acid (mobile phase B). The following 12.5 minute gradient was used: 1) 0-1 min, 5% B; 2) 1-8 min, linear increase to 100% B; 3) 8-10 min, 100% B; 4) 10-10.5 min, linear decrease to 5% B; 5) 10.5-12 min, 5% B. A flow rate of 0.5 mL/min was maintained, and the column chamber temperature was kept at 40°C.

For MS/MS acquisition, a Q Exactive Plus (Thermo Scientific) high-resolution mass spectrometer was used. Both positive and negative mode data were collected, with the parameters shown in **(Supplementary Table 12)**. Data acquisition was performed in randomized order. Injection volume was 5 μL. A blank and pooled quality control samples were analyzed every 12 injections to monitor instrument performance. Retention time shifts were analyzed by using a mix of sulfadimethoxine, sulfachloropyridazine, sulfamethazine, sulfamethizole, amitriptyline, and coumarin-314 standards. This standard mix was taken at the beginning of the run, every 100 samples throughout the run, and at the end of the run.

The Q Exactive Plus was calibrated using Pierce LTQ Velos ESI positive and negative ion calibration solution (ThermoFisher) prior to analysis.

### Data Processing

Raw files were converted to mzML files using MSConvert ^27^. Classical Molecular Networking was performed using GNPS ^5^. The parameters used are listed in **Supplementary Table 9**, and we found that varying the parameters did not significantly impact the rational library building (**Supplementary Data 1**). Molecular networks were visualized using Cytoscape version 3.9.1. All data processing was done in Rstudio version 4.3.0 or Jupyter Notebook 6.5.4. The map of soil sample collection locations was done in qGIS 3.30.3.

Figure 1A generated using BioRender.com

Rational Libraries were generated using an R studio code made freely available (See Data Availability Section). This code uses the node table from the Classical Molecular networking job, which summarizes each feature, its scaffold family, and which extracts contain the feature. To choose extracts to be added to the rational library, it aggregates the features by scaffold, then chooses the extract that contains the most scaffold . Then, those scaffolds are deleted from the dataset. Then, the extract with the most scaffolds not already accounted for is added into the rational library. This process repeats until a desired percent of maximum diversity is reached.

Bioactivity correlations were done in R studio with code that is also freely available, loosely adapted from the method described by ^19^. This code requires the Classical Molecular Networking Bucket Table, which summarizes each feature and its sum precursor abundance within each sample. We used this as an approximate indication of relative abundance. The code requires the input of the activity data for one of the assays. Then, it builds a Pearson correlation (Regular or Pearson’s Bivariate Correlation Coefficient, depending on if the activity data is continuous or binary) between the number of scans and the activity. It also calculates and corrects the p-value of each feature to activity. The features are filtered based on which ones have a p-value less than 0.05, and a Pearson Correlation statistic greater than 0 (positively correlated with activity).

### Data Availability

The .raw and .mzML data files used in this article are available in the MassIVE repository under accession numbers MSV000091950 (positive data), and MSV000091980 (negative data). The Classical Molecular Networking jobs are available at:

https://gnps.ucsd.edu/ProteoSAFe/status.jsp?task=37a75c380d7c464ba278433c6434f7c1 (Fungal extracts, positive data, parameters listed in Supplementary Table 9)

https://gnps.ucsd.edu/ProteoSAFe/status.jsp?task=ab2cceee30e347d185e17d3e2dc95092 (Fungal extracts, negative data, parameters listed in Supplementary Table 9)

https://gnps.ucsd.edu/ProteoSAFe/status.jsp?task=8ad5a0af7cbc4859a7b943fcaebb7e9d (Low resolution mimicking data, Fungal extracts, Positive data, parameters listed in Supplementary Table 9). The jobs used to verify parameter sensitivity are listed in (Supplementary Data 1)

The publicly available data^12^ used to verify our method is available in the Massive repository under accession number MSV000087728. The Classical Molecular Networking job is available at https://gnps.ucsd.edu/ProteoSAFe/status.jsp?task=a0349e711a214913b0222041e76c2346.

The codes used, including for library building and bioactivity correlations are available at https://github.com/mmness/. The bioactivity correlation code is loosely based on the technique presented by Nothias et al ^19^.

The Genbank accession numbers used for sequencing are PP664564 - PP665462.

## Acknowledgements

This project was supported by NIH awards R01GM145649 and R01AI154777. The content is solely the responsibility of the authors and does not necessarily represent the official views of the National Institutes of Health.

## References

1. Newman, D. J. & Cragg, G. M. Natural Products as Sources of New Drugs from 1981 to 2014. J. Nat. Prod. 79, 629–661 (2016).

2. Clark, C. M., Nguyen, L., Pham, V. C., Sanchez, L. M. & Murphy, B. T. Automated Microbial Library Generation Using the Bioinformatics Platform IDBac. Molecules 27, (2022).

3. Anderson, V. M., Wendt, K. L., Najar, F. Z.McCall, L.-I. & Cichewicz, R. H. Building Natural Product Libraries Using Quantitative Clade-Based and Chemical Clustering Strategies. mSystems 6, (2021).

4. Costa, M. S., Clark, C. M., Ómarsdóttir, S., Sanchez, L. M. & Murphy, B. T. Minimizing Taxonomic and Natural Product Redundancy in Microbial Libraries using MALDI-TOF MS and the bioinformatics pipeline IDBac. J. Nat. Prod. 82, 2167 (2019).

5. Wang, M. et al. Sharing and community curation of mass spectrometry data with Global Natural Products Social Molecular Networking. Nat. Biotechnol. 34, 828–837 (2016).

6. Hernandez, A., Nguyen, L. T., Dhakal, R. & Murphy, B. T. The need to innovate sample collection and library generation in microbial drug discovery: a focus on academia. Nat. Prod. Rep. 38, 292–300 (2021).

7. Kim, S. et al. PubChem structure-activity relationship (SAR) clusters. J. Cheminform. 7, 33 (2015).

8. Lassalas, P. et al. Structure Property Relationships of Carboxylic Acid Isosteres. J. Med. Chem. 59, 3183–3203 (2016).

9. Pinheiro, P. de S. M., Franco, L. S. & Fraga, C. A. M. The Magic Methyl and Its Tricks in Drug Discovery and Development. Pharmaceuticals 16, (2023).

10. Hughes, J. P., Rees, S., Kalindjian, S. B. & Philpott, K. L. Principles of early drug discovery. Br. J. Pharmacol. 162, 1239–1249 (2011).

11. Crüsemann, M. et al. Prioritizing Natural Product Diversity in a Collection of 146 Bacterial Strains Based on Growth and Extraction Protocols. J. Nat. Prod. 80, 588–597 (2017).

12. Allard, P.-M. et al. Open and reusable annotated mass spectrometry dataset of a chemodiverse collection of 1,600 plant extracts. Gigascience 12, giac124 (2023).

13. Smith, C. A. et al. METLIN: a metabolite mass spectral database. Ther. Drug Monit. 27, 747–751 (2005).

14. Website. Dictionary of Natural Products, (a.n.d.). https://dnp.chemnetbase.com/chemical/ChemicalSearch.xhtml?dswid=-6075.

15. Website. NIST Standard Reference Database, (n.d.). https://www.nist.gov/srd.

16. Qin, G.-F. et al. MS/MS-Based Molecular Networking: An Efficient Approach for Natural Products Dereplication. Molecules 28, (2022).

17. Natural Products Atlas. https://www.npatlas.org/discover/snapms/.

18. Natural Products Atlas. https://www.npatlas.org/discover/snapms/.

19. Nothias, L.-F. et al. Bioactivity-Based Molecular Networking for the Discovery of Drug Leads in Natural Product Bioassay-Guided Fractionation. J. Nat. Prod. 81, 758–767 (2018).

20. Hossain, E. et al. Mapping of host-parasite-microbiome interactions reveals metabolic determinants of tropism and tolerance in Chagas disease. Sci Adv 6, eaaz2015 (2020).

21. Du, L. et al. Crowdsourcing natural products discovery to access uncharted dimensions of fungal metabolite diversity. Angew. Chem. Int. Ed Engl. 53, 804–809 (2014).

22. King, J. B. et al. Design and Application of a High-Throughput, High-Content Screening System for Natural Product Inhibitors of the Human Parasite. ACS Infect Dis 5, 1456–1470 (2019).

23. Du, L. et al. Diarylcyclopentendione metabolite obtained from a Preussia typharum isolate procured using an unconventional cultivation approach. J. Nat. Prod. 75, 1819–1823 (2012).

24. Anderson, V. M. et al. Assessing Microbial Metabolic and Biological Diversity to Inform Natural Product Library Assembly. J. Nat. Prod. 85, 1079–1088 (2022).

25. Collins, J. E. et al. Antiplasmodial peptaibols act through membrane directed mechanisms. Cell Chem Biol 31, 312–325.e9 (2024).

26. Trager, W. & Jensen, J. B. Human malaria parasites in continuous culture. Science 193, 673–675 (1976).

27. Chambers, M. C. et al. A cross-platform toolkit for mass spectrometry and proteomics. Nat. Biotechnol. 30, 918–920 (2012).

