## Supplemental figures and tables for "Rationally Minimizing Natural Product Libraries Using Mass Spectrometry"

**Supplementary Information for Rationally Minimizing Natural Product Libraries Using Mass Spectrometry**

This pdf includes:

__Supplemental Tables 1-12__

**Supplemental Table 1:** Summary of rational library sizes with different parameters for scaffold building.

**Supplemental Table 2:** Hit rates for random sampling of positive mode fungal data.

**Supplemental Table 3:** Retention of features correlated to activity in the rational libraries, for positive mode analysis of fungal samples.

**Supplemental Table 4:** Summary of rational library sizes using publicly available data.

**Supplemental Table 5:** Higher hit rates for rational libraries, using publicly available data.

**Supplemental Table 6:** Retention of features correlated to activity in the rational libraries, in publicly available data.

**Supplemental Table 7:** Higher hit rates for rational libraries, for negative mode analysis of fungal samples.

**Supplemental Table 8:** Retention of features correlated to activity in the rational libraries, using negative mode analysis of fungal samples.

**Supplemental Table 9:** Classical molecular networking parameters.

**Supplemental Table 10:** Higher hit rates for rational libraries, using low-resolution mimicking parameters and positive mode analysis of fungal samples.

**Supplemental Table 11:** Retention of features correlated to activity in the rational libraries, using low-resolution mimicking parameters and positive mode analysis of fungal samples.

**Supplemental Table 12:** Q-Exactive Plus parameters used for fungal data collection.

__Supplemental Figures 1-7__

**Supplemental Figure 1:** Rapid accumulation of scaffold diversity with our rational library building method, applied to publicly available data.

**Supplemental Figure 2:** Retention of features correlated to activity in the rational libraries, on publicly available data.

**Supplemental Figure 3:** Rapid accumulation of scaffold diversity with our rational library building method, for negative mode analysis of fungal samples.

**Supplemental Figure 4:** Retention of features correlated to activity in the rational libraries, for negative mode analysis of fungal samples.

**Supplemental Figure 5:** Rapid accumulation of scaffold diversity with our rational library building method, using low-resolution mimicking parameters and positive mode analysis of fungal samples.

**Supplemental Figure 6:** Retention of features correlated to activity in the rational libraries, using low-resolution mimicking parameters and positive mode analysis of fungal samples.

**Supplemental Figure 7:** Fungal samples collection sites.

**Supplemental Tables:**

**Supplemental Table 1: Summary of rational library sizes with different parameters for scaffold building.** Both positive and negative mode data were collected. Applying our method to the negative mode data, and to low resolution-mimicking data (based on data processing parameters) resulted in similar reduction effectiveness with regards to library size reduction.

| | Number of Extracts to 80% Maximum Scaffold diversity | Number of Extracts to 95% Maximum Scaffold diversity | Number of Extracts to 100% Maximum Scaffold diversity |
|---|---|---|---|
| Positive Mode Fungal Data | 50 (96.5% reduction) | 116 (91.9% reduction) | 216 (84.9% reduction) |
| Negative Mode Fungal Data | 41 (97.1% reduction) | 96 (93.3% reduction) | 174 (87.9% reduction) |
| Positive Mode Fungal Data: Low Resolution Mimicking Data | 44 (96.9% reduction) | 106 (92.6% reduction) | 195 (86.4% reduction) |

**Supplemental Table 2: Hit rates for random sampling of positive mode fungal data.** To prove our method's increased hit rates are not an artifact of a reduced absolute library size, we created 1000 iterations of random sampling that were the same size as our rationally reduced library (see Supplemental Table 1, positive fungal mode). Hit rates against each activity assay were then calculated. The lower quartiles, medians, and upper quartiles for all assay hit rates are listed. In every case, our rational method (Table 1) outperforms the random sampling.

| Activity assay | Lower quartile, median, and upper quartile hit rates for 50 random extracts (1000 iterations) | Lower quartile, median, and upper quartile hit rates for 116 random extracts (1000 iterations) | Lower quartile, median, and upper quartile hit rates for 216 random extracts (1000 iterations) |
|---|---|---|---|
| *P. falciparum* | 8.00%<br>12.00%<br>14.00% | 9.48%<br>11.21%<br>12.93% | 9.72%<br>11.11%<br>12.50% |
| *T. vaginalis* | 4.00%<br>8.00%<br>10.00% | 6.02%<br>7.76%<br>9.04% | 6.48%<br>7.87%<br>8.80% |
| Neuraminidase | 0.00%<br>2.00%<br>2.00% | 0.86%<br>1.72%<br>2.58% | 1.39%<br>1.85%<br>2.31% |

**Supplemental Table 3: Retention of features correlated to activity in the rational libraries, for positive mode analysis of fungal samples.** The correlation between each feature detected by the LC-MS/MS run and the activity was calculated. The significance cutoff was a Bonferroni corrected p-value of $p < 0.05$, and a Pearson correlation of $r > 0$. After identifying the significantly correlated features, their retention in the rational libraries was assessed. Visualization of the data is shown in **Figure 1c.**

| Activity assay | Features found to be significantly correlated to activity in full dataset | Retained in the 80% scaffold diversity library | Retained in the 95% scaffold diversity library | Retained in the 100% scaffold diversity library |
|---|---|---|---|---|
| *P. falciparum* | 266 | 223 (83.8%) | 247 (92.9%) | 260 (97.7%) |
| *T. vaginalis* | 205 | 179 (87.3) | 195 (95.1%) | 204 (99.5%) |
| Neuraminidase | 125 | 92 (73.6%) | 104 (83.2%) | 105 (84.0%) |

**Supplemental Table 4: Summary of rational library sizes using publicly available data.** Data was obtained from reference [12]**,** as deposited in the MassIVE repository under accession number MSV000087728.This collection consisted of a broad taxonomy of plant exacts.

| | Number of Extracts to 80% Maximum Scaffold diversity | Number of Extracts to 95% Maximum Scaffold diversity | Number of Extracts to 100% Maximum Scaffold diversity |
|---|---|---|---|
| Public Data (1600 total) | 104 | 233 | 408 |

**Supplemental Table 5: Higher hit rates for rational libraries, using publicly available data.** Reference [12], which described LC-MS data for plant extracts, also included bioactivity assay against *Trypanosoma cruzi* parasites, causative agents of Chagas disease. We therefore assessed hit rates using the rational libraries selected as described in **Supplemental Table 4**. Results of the hit rates in the rational libraries compared to the full library are summarized.

| Activity assay | Hit Rate in full Library | Hit rate in the 80% scaffold diversity library | Hit rate in the 95% scaffold diversity library | Hit rate in the 100% scaffold diversity library |
|---|---|---|---|---|
| *T. cruzi* | 0.5% | 1.92% | 1.72% | 1.47% |

**Supplemental Table 6: Retention of features correlated to activity in the rational libraries, in publicly available data.** The correlation between each feature detected by LC-MS/MS in reference [12] and the reported anti-*T. cruzi* activity was calculated (See Data Availability Section). The significance cutoff was a FDR corrected p-value of $p < 0.05$, and a Pearson's Bivariate Correlation Coefficient correlation of $r > 0$. This correlation method was selected because *T. cruzi* activity was provided as binomial data rather than continuous data. After identifying the significantly correlated features, their retention in the rational libraries were confirmed. Visualization of this data is in **Supplemental Fig. 2**

| Activity assay | Features found to be significantly correlated to activity in full dataset | Retained in the 80% scaffold diversity library | Retained in the 95% scaffold diversity library | Retained in the 100% scaffold diversity library |
|---|---|---|---|---|
| *T. cruzi* | 108 | 84<br>(77.8%) | 95<br>(88.0%) | 103<br>(95.4%) |

**Supplemental Table 7. Higher hit rates for rational libraries, for negative mode analysis of fungal samples.** We assessed whether building scaffolds and the rational libraries based on the negative polarity MS/MS data yielded the same results as analysis of data collected in positive mode. We found similar patterns, with an increased hit rate in the rational libraries for each activity assay.

| Activity assay | Hit Rate in full Library | Hit rate in the 80% scaffold diversity library | Hit rate in the 95% scaffold diversity library | Hit rate in the 100% scaffold diversity library |
|---|---|---|---|---|
| *P. falciparum* | 11.26% | 26.83% | 18.75% | 13.22% |
| *T. vaginalis* | 7.64% | 12.20% | 12.50% | 9.77% |
| Neuraminidase | 2.57% | 7.32% | 4.17% | 3.45% |

**Supplemental Table 8: Retention of features correlated to activity in the rational libraries, using negative mode analysis of fungal samples.** The correlation between each feature detected by the negative polarity LC-MS/MS run and the activity was calculated (See Data Availability Section). The significance cutoff was a Bonferroni corrected p-value of $p < 0.05$, and a Pearson correlation of $r > 0$. After identifying the significantly correlated features, their retention in the rational libraries were confirmed. Visualization of the data is in **Supplemental Fig. 4.**

| Activity assay | Features found to be significantly correlated to activity in full dataset | Retained in the 80% scaffold diversity library | Retained in the 95% scaffold diversity library | Retained in the 100% scaffold diversity library |
|---|---|---|---|---|
| *P. falciparum* | 103 | 80 (77.7%) | 87 (84.5%) | 97 (94.2%) |
| *T. vaginalis* | 117 | 102 (87.2%) | 105 (89.7%) | 107 (91.5%) |
| Neuraminidase | 134 | 99 (73.9%) | 101 (75.4%) | 114 (85.1%) |

**Supplemental Table 9: Classical molecular networking parameters**. Parameters used for the positive polarity fungal data, public data, and negative polarity fungal data ("Analysis Parameters"). For low-resolution mimicking parameters, only positive polarity fungal samples were analyzed. See Data Availability section for GNPS job links.

| | Analysis Parameters | Low-Resolution Mimicking Parameters |
|---|---|---|
| **Basic Options** | | |
| Precursor Ion Mass Tolerance | 0.02 Da | 2 Da |
| Precursor Ion Mass Tolerance | 0.02 Da | 0.95 Da |
| **Advanced Network Options** | | |
| Minimum Pairs Cosine | 0.7 | 0.7 |
| Network TopK | 7 | 7 |
| Maximum Connected Component Size | 70 | 70 |
| Minimum Matched Fragment Ions | 4 | 4 |
| Minimum Cluster Size | 4 | 4 |
| Maximum shift | 500 Da | 500 Da |
| **Advanced Filtering Options** | | |
| Filter below Standard Deviation | 0 | 0 |
| Filter Precursor Window | filter | filter |
| Filter peaks in 50 Da Window | filter | filter |
| Filter Spectra from G6 as Blanks Before Networking | don't filter | don't filter |
| Minimum Peak Intensity | 0 | 0 |
| Filter Library | filter library | filter library |

**Supplemental Table 10: Higher hit rates for rational libraries, using low-resolution mimicking parameters and positive mode analysis of fungal samples.** To the same positive polarity fungal extract data, classical molecular networking was performed with parameters recommended for low-resolution LC-MS/MS data (See **Supplemental Table 9.** for parameters). The scaffolds and rational libraries were built using this output. We found similar patterns between the low-resolution mimicking and regular parameters, evident by an increased hit rate in the rational libraries for each activity assay.

| Activity assay | Hit Rate in full Library | Hit rate in the 80% scaffold diversity library | Hit rate in the 95% scaffold diversity library | Hit rate in the 100% scaffold diversity library |
|---|---|---|---|---|
| *P. falciparum* | 11.26% | 13.64% | 12.26% | 12.31% |
| *T. vaginalis* | 7.64% | 15.91% | 10.38% | 9.23% |
| Neuraminidase | 2.57% | 4.55% | 3.77% | 4.62% |

**Supplemental Table 11: Retention of features correlated to activity in the rational libraries, using low-resolution mimicking parameters and positive mode analysis of fungal samples.** The correlation between each feature detected by the LC-MS/MS run and the activity was calculated. The significance cutoff was a Bonferroni corrected p-value of $p < 0.05$, and a Pearson correlation of $r > 0$. After identifying the significantly correlated features, their retention in the rational libraries was confirmed. Visualization of the data is in **Supplemental Fig. 6**.

| Activity assay | Features found to be significantly correlated to activity in full dataset | Retained in the 80% scaffold diversity library | Retained in the 95% scaffold diversity library | Retained in the 100% scaffold diversity library |
|---|---|---|---|---|
| *P. falciparum* | 199 | 150 (75.4%) | 169 (84.9%) | 187 (94.0%) |
| *T. vaginalis* | 140 | 102 (72.9%) | 117 (83.6%) | 132 (94.3%) |
| Neuraminidase | 105 | 87 (82.9%) | 93 (88.6%) | 103 (98.1) |

**Supplemental Table 12. Q-Exactive Plus parameters used for fungal data collection.** Summary of parameters used in the Thermo Scientific XCalibur software for the positive and negative fungal extract data collection.

| Parameter | Value (positive polarity) | Value (negative polarity) |
|---|---|---|
| Default Charge State | 1 | 1 |
| Polarity | Positive | Negative |
| Runtime | 12.5 min | 12.5 min |
| **Tune data** | | |
| Ion source | HESI | HESI |
| Capillary temp (°C) | 320 | 320 |
| Spray voltage (V) | 3800 | 3000 |
| Sheath gas (Thermo Arbitrary units) | 35 | 35 |
| Max spray current (μA) | 100 | 100 |
| Probe heater temp (°C) | 350 | 350 |
| S-lens Radiofrequency level (Thermo Arbitrary units) | 50 | 50 |
| Auxiliary gas (Thermo Arbitrary units) | 10 | 10 |
| Sweep gas (Thermo Arbitrary units) | 0 | 0 |
| **Full MS** | | |
| Scan Range | 100-1500 *m/z* | 100-1500 *m/z* |
| Maximum Injection Time | 246 ms | 246 ms |
| Resolution (at *m/z* 200, full width at half maximum (FWHM)) | 70,000 | 70,000 |
| AGC Target | 1e6 | 1e6 |
| **dd-MS2** | | |
| Isolation Window | 1.0 *m/z* | 1.0 *m/z* |
| Maximum Injection Time | 54 ms | 54 ms |

| | | |
|---|---|---|
| (N)CE/stepped(N)CE | 20 ,40, 60 | 20, 40, 60 |
| Resolution (at *m/z* 200, full width at half maximum (FWHM)) | 17,500 | 17,500 |
| AGC Target | 2e5 | 2e5 |
| TopN | 5 | 5 |
| **dd Settings** | | |
| Intensity threshold | 1.5e5 | 1.5e5 |
| Minimum AGC target | 8.00e3 | 8.00e3 |
| Dynamic exclusion | 10 s | 10 s |
| Exclude isotopes | on | on |
| Peptide match | preferred | preferred |
| AGC: Automatic Gain Control, N(CE): Normalized Collision Energy, HESI: Heated electrospray ionization | | |

**Supplemental Figures:**

**Supplemental Figure 1: Rapid accumulation of scaffold diversity with our rational library building method, applied to publicly available data.** Scaffold accumulation for random sample selection (50 iterations, each until 100% scaffold diversity reached), outperformed by our rational library selection method.

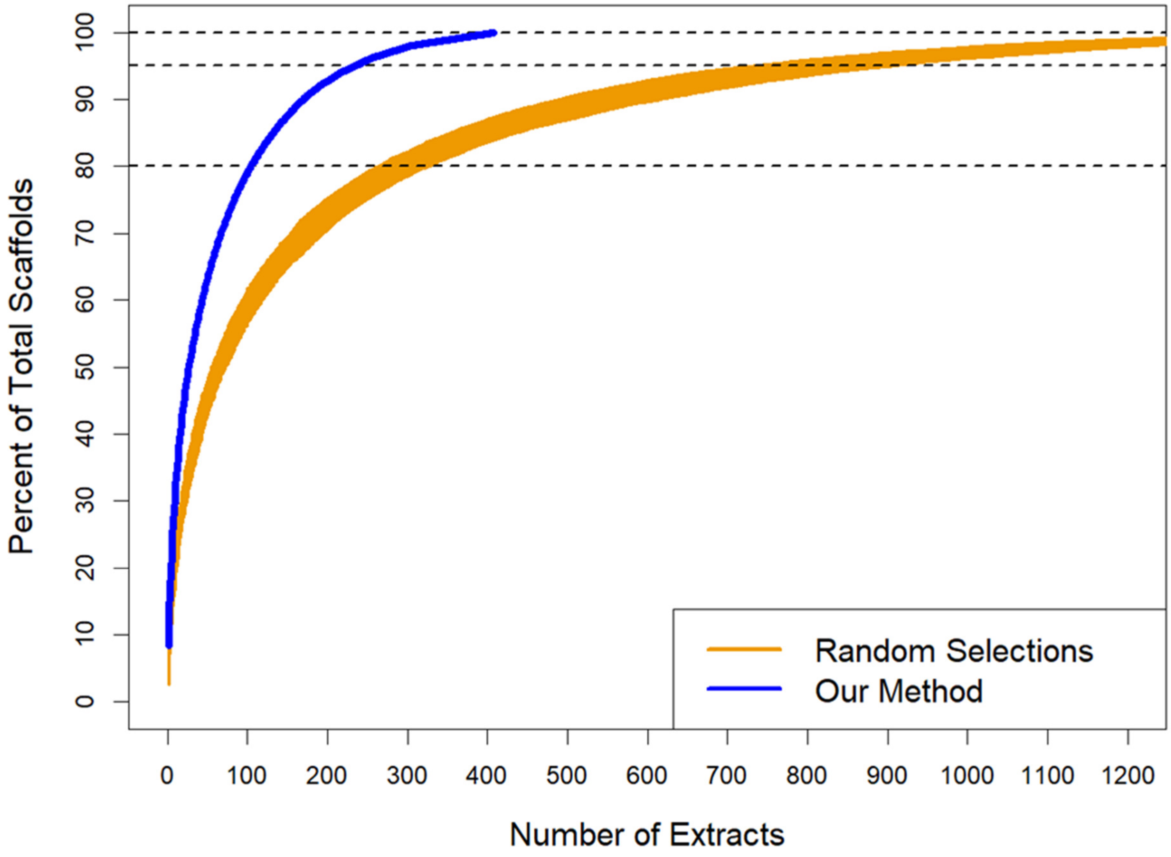

**Supplemental Figure 2: Retention of features correlated to activity in the rational libraries, on publicly available data.** The correlation between each feature abundance, determined by LC-MS/MS, and anti-*T. cruzi* activity in publicly-available data was calculated (See Data Availability Section). The significance cutoff was a FDR corrected p-value of $p < 0.05$, and a Pearson's Bivariate Correlation Coefficient correlation of $r > 0$. This correlation method was selected because *T. cruzi* activity was published as binomial data rather than continuous data. After identifying the significantly correlated features, their retention in the rational libraries was confirmed. Table of numerical data is shown in **Supplemental Table 6.**

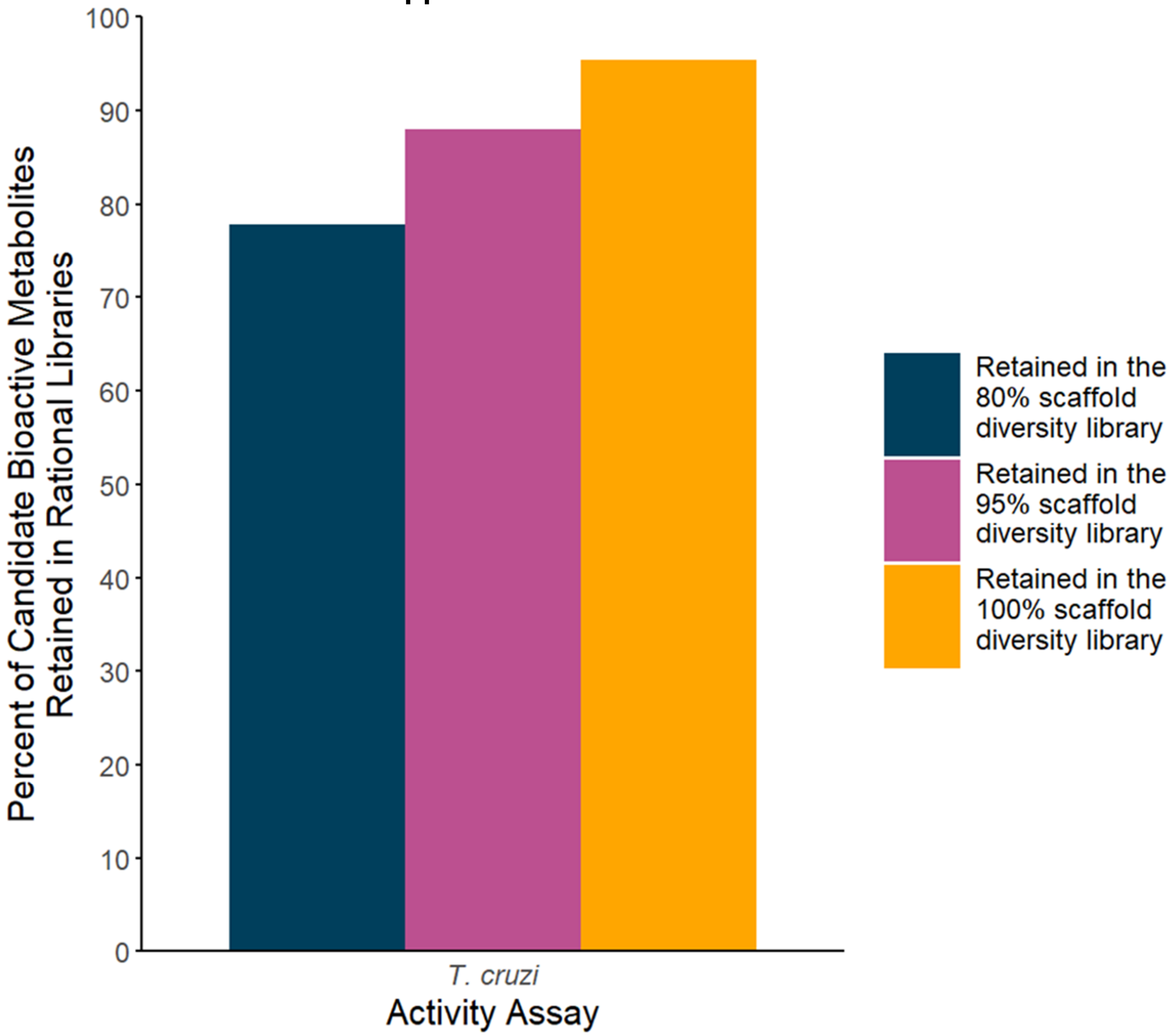

**Supplemental Figure 3: Rapid accumulation of scaffold diversity with our rational library building method, for negative mode analysis of fungal samples.** Scaffold accumulation for random sample iterations (50 iterations, each until 100% scaffold diversity reached), outperformed by our rational library selection method.

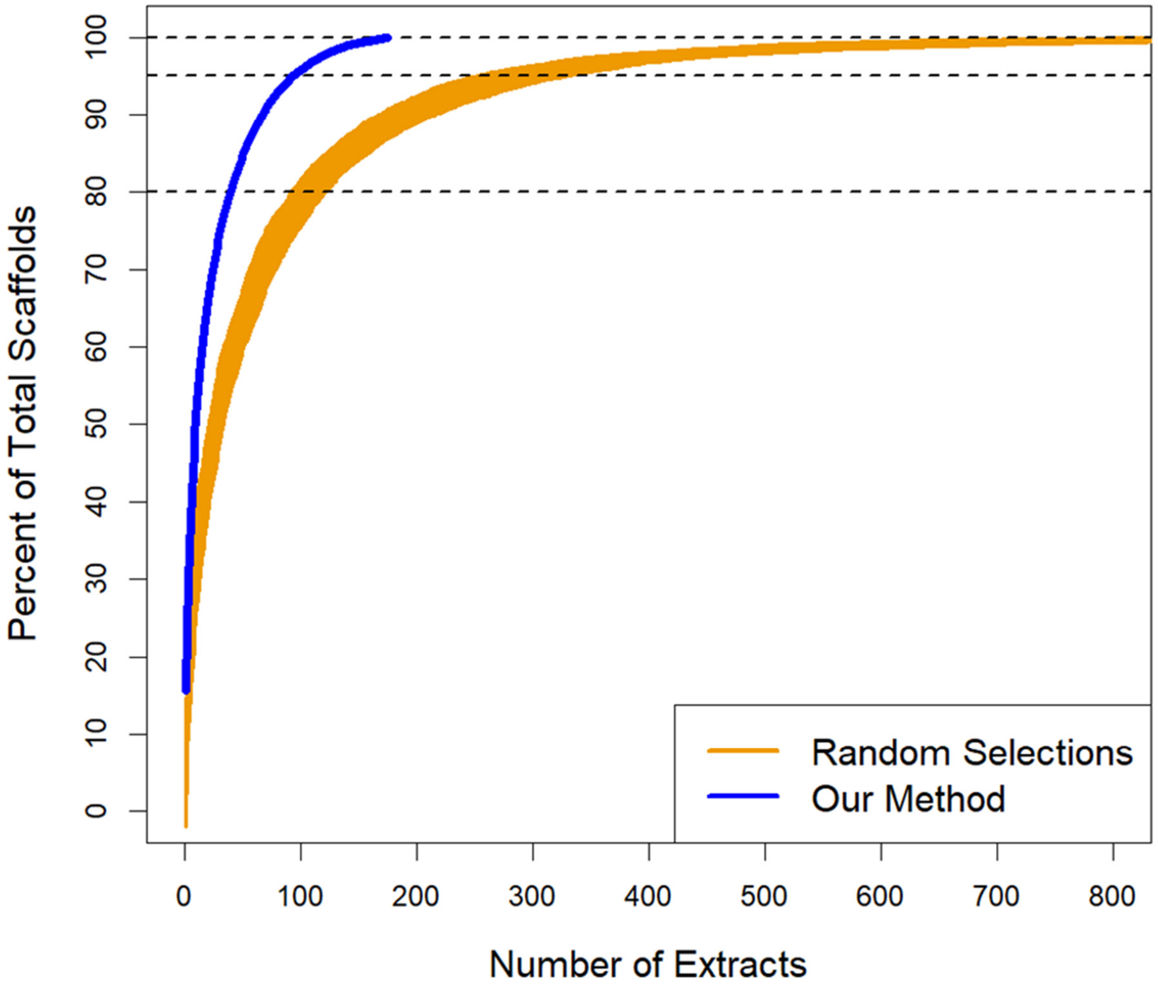

**Supplemental Figure 4: Retention of features correlated to activity in the rational libraries, for negative mode analysis of fungal samples.** The correlation between the presence of each feature detected by the negative polarity LC-MS/MS run and activity was calculated (See Data Availability Section). The significance cutoff was a Bonferroni corrected p-value of $p < 0.05$, and a Pearson correlation of $r > 0$. After identifying the significantly correlated features, their retention in the rational libraries was confirmed. Table of numerical data is shown in **Supplemental Table 8.**

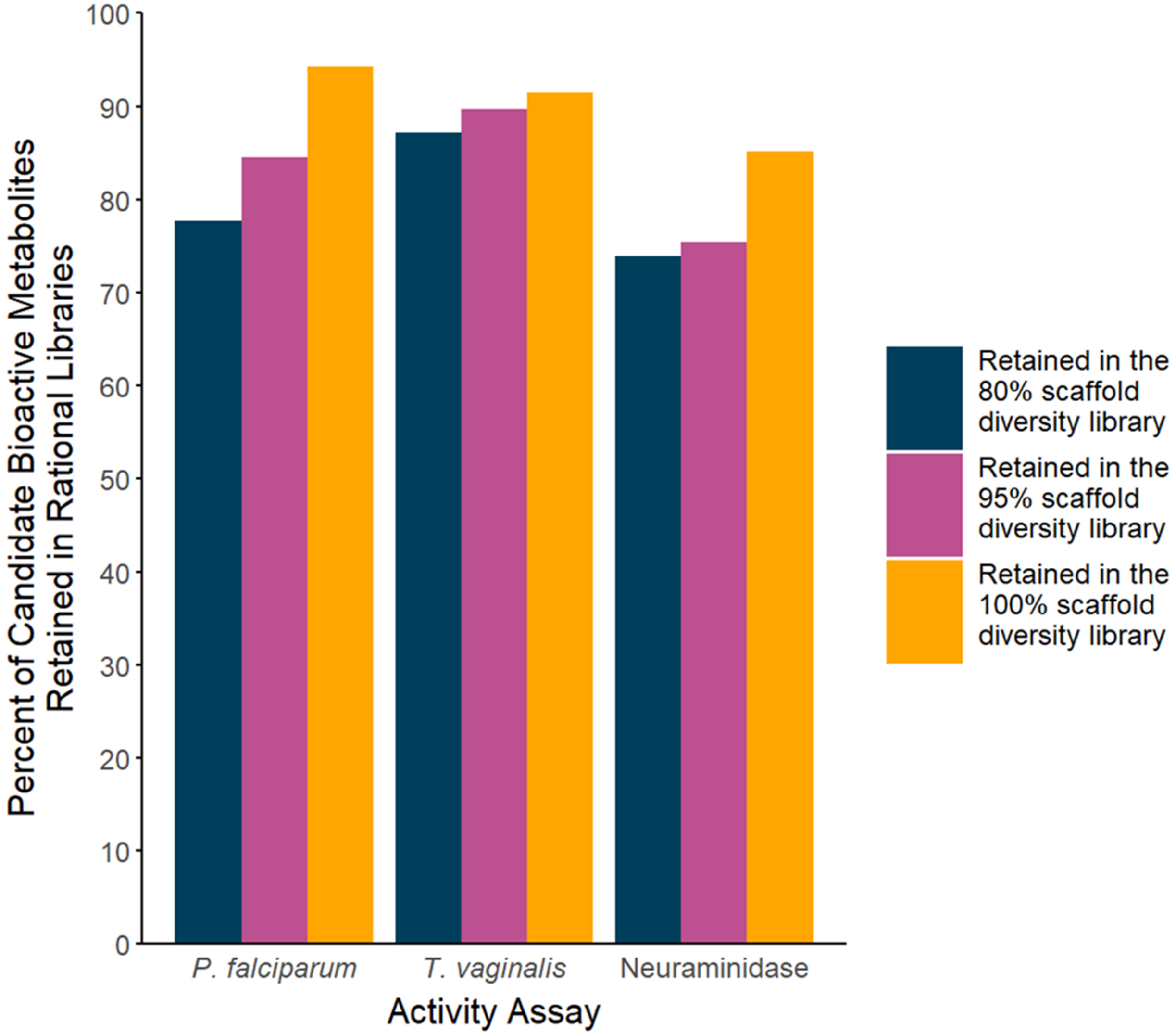

**Supplemental Figure 5: Rapid accumulation of scaffold diversity with our rational library building method, using low-resolution mimicking parameters and positive mode analysis of fungal samples.** Scaffold accumulation of random sample iterations (50 iterations, each until 100% scaffold diversity reached), compared to our rational library selection method applied on data processed with low resolution-mimicking parameters.

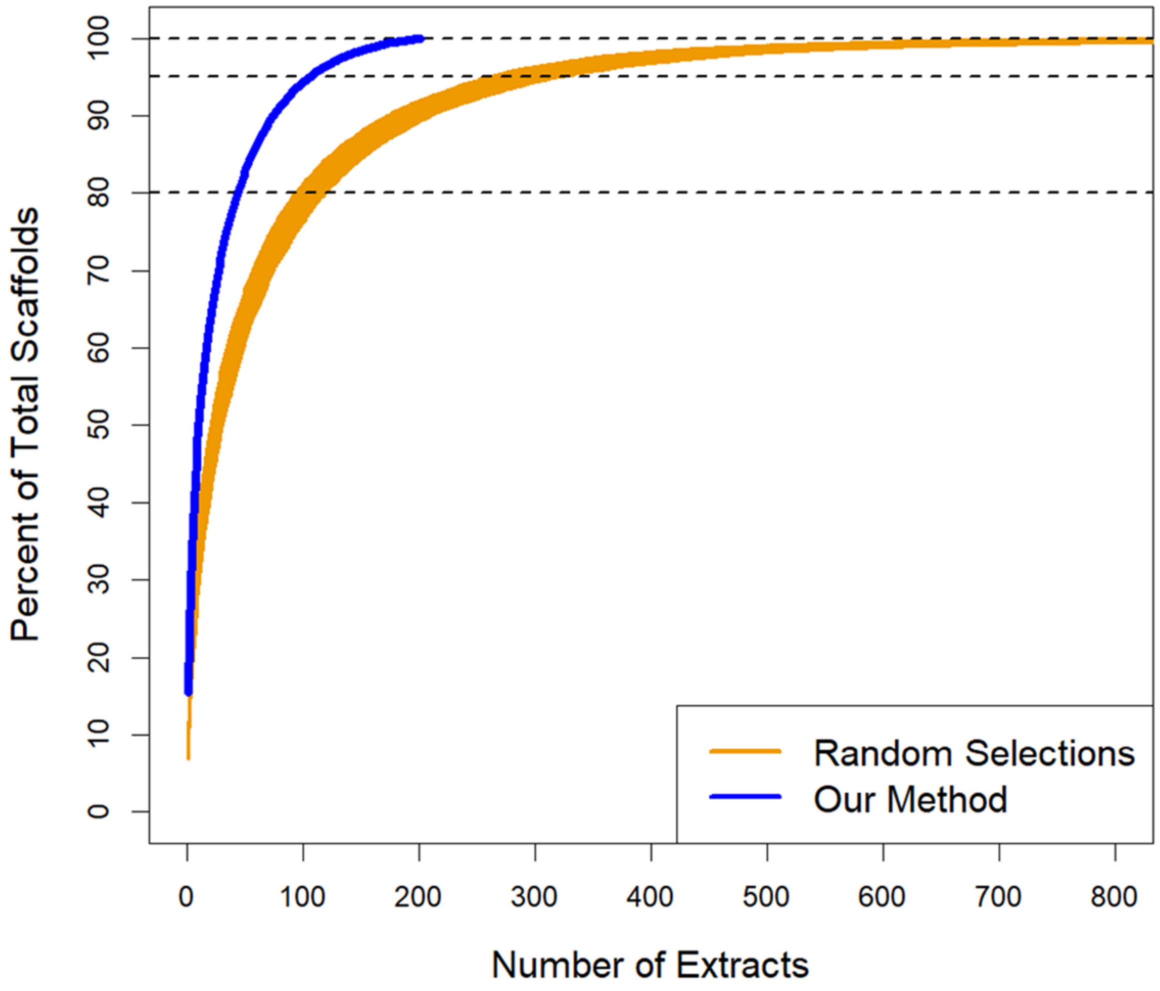

**Supplemental Figure 6: Retention of features correlated to activity in the rational libraries, using low-resolution mimicking parameters and positive mode analysis of fungal samples.** The correlation between the abundance of each feature detected by the LC-MS/MS run and activity was calculated (See Data Availability Section). The significance cutoff was a Bonferroni corrected p-value of $p < 0.05$, and a Pearson correlation of $r > 0$. After identifying the significantly correlated features, their retention in the rational libraries were confirmed. Table of numerical data is shown in **Supplemental Table 12.**

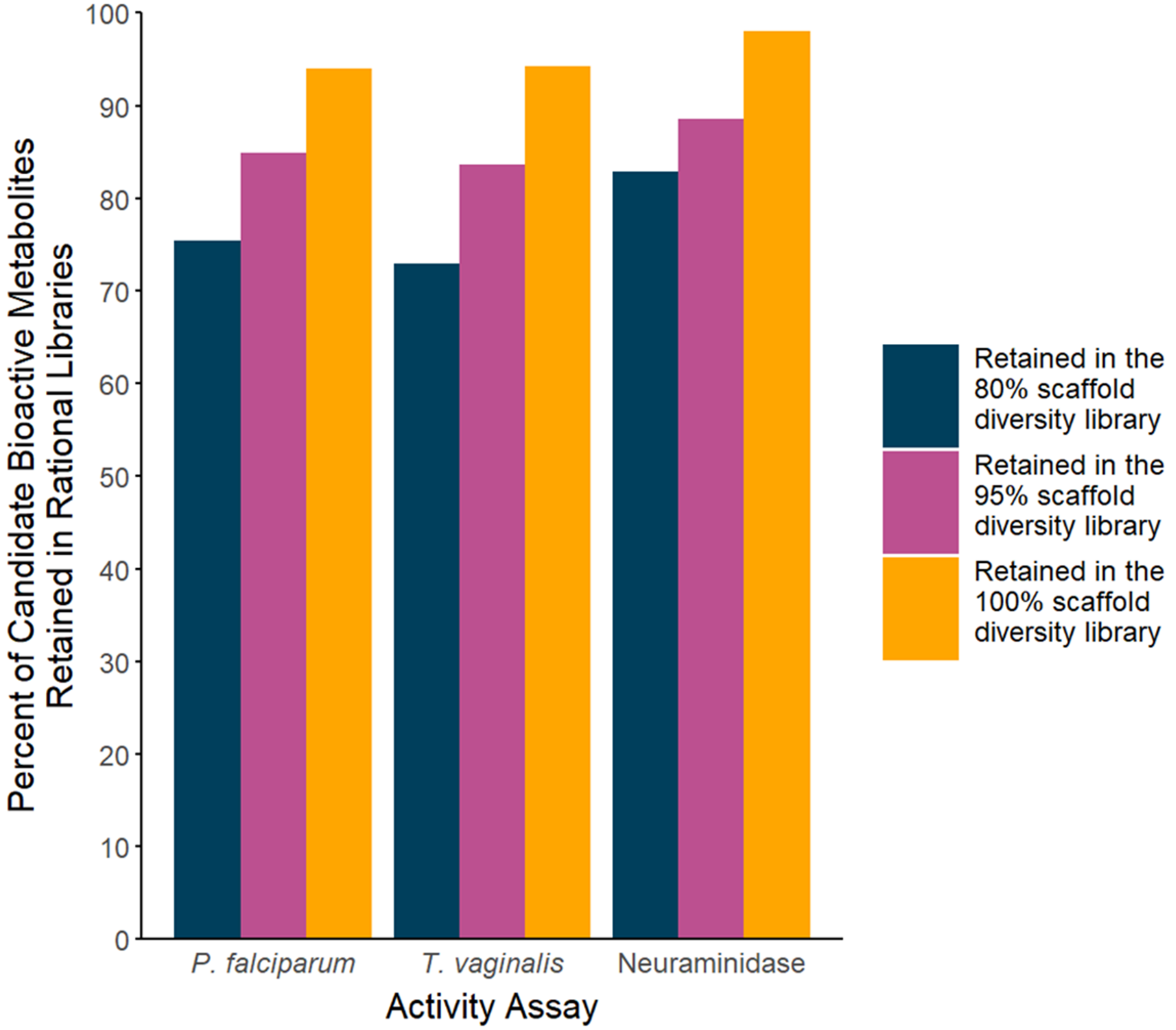

**Supplemental Figure 7**: **Fungal samples collection sites.** Summary of the origins of the fungal extracts used for analysis.

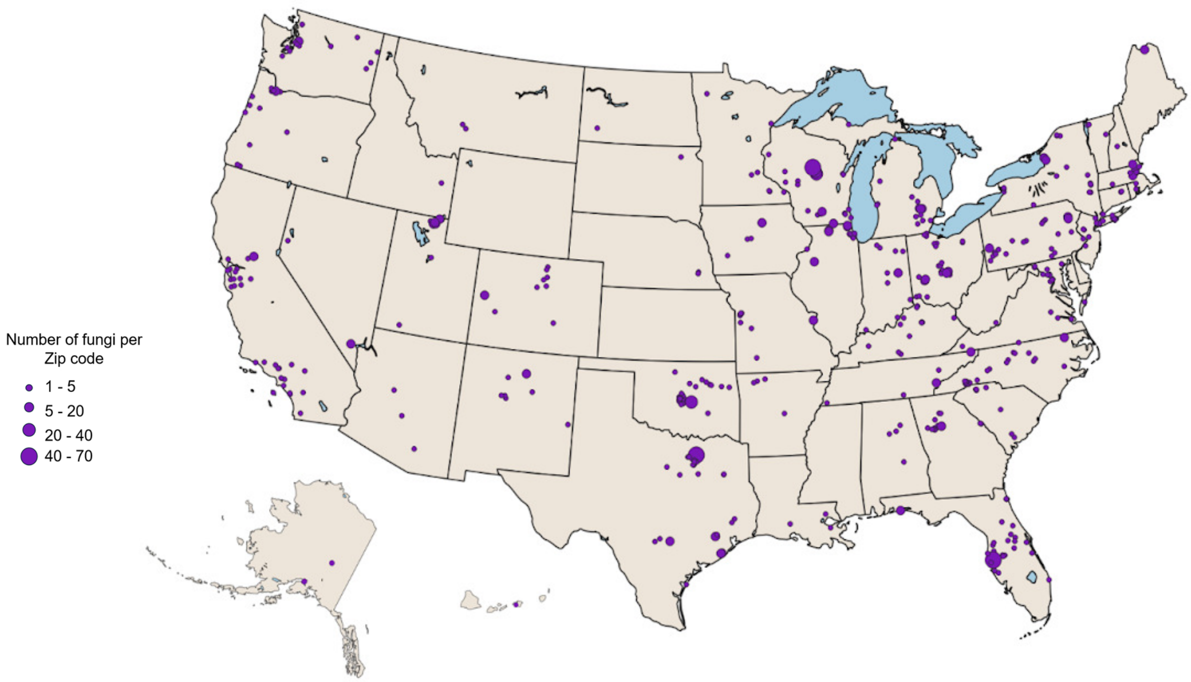

**Supplementary Data:**

**Supplemental Data Sheet 1: Changing classical molecular networking parameters has little impact on library size and hit rates.** Summary of hit rates and library sizes with the varying of multiple molecular networking parameters. Note that other than the parameters listed in the table, all other parameters are the same as listed in **Supplemental Table 9.**

**Supplemental Data Sheet 2: Fungal Metadata.** Summary of each fungal extract, the zip code the sample was collected, and the genus.
